# Efficient targeted transgenesis of large donor DNA into multiple mouse genetic backgrounds using bacteriophage Bxb1 integrase

**DOI:** 10.1101/2021.09.20.461117

**Authors:** Benjamin E. Low, Vishnu Hosur, Simon Lesbirel, Michael V. Wiles

## Abstract

Efficient, targeted integration of large DNA constructs represent a significant hurdle in genetic engineering for the development of mouse models of human disease and synthetic biology research. To address this, we developed a system for efficient and precise, targeted single-copy integration of large transgenes directly into the zygote using multiple mouse genetic backgrounds. Conventional approaches, such as random transgenesis, CRISPR/Cas9-mediated homology-directed repair (HDR), lentivirus-based insertion, or DNA transposases all have significant limitations. Our strategy uses *in vivo* Bxb1 mediated recombinase-mediated cassette exchange (RMCE) to efficiently generate precise single-copy integrations of transgenes. This is achieved using a transgene “landing pad” composed of dual heterologous Bxb1 attachment (att) sites in *cis*, pre-positioned in the *Gt(ROSA)26Sor* safe harbor locus. Successful RMCE is achieved in att carrier zygotes using donor DNA carrying cognate attachment sites flanking the desired donor transgene microinjected along with Bxb1-integrase mRNA. This approach routinely achieves perfect vector-free integration of donor constructs at efficiencies as high as 43% and has generated transgenic animals containing inserts up to ∼43kb. Furthermore, when coupled with a nanopore-based Cas9-targeted sequencing (nCATS) approach, complete verification of the precise insertion sequence can be achieved. As a proof-of-concept we describe the creation and characterization of C57BL/6J and NSG Krt18-ACE2 transgenic mouse models for SARS-CoV2 research with verified heterozygous N1 animals available for experimental use in ∼4 months. In addition, we created a diverse series of mouse backgrounds carrying a single att site version of the landing pad allele in C57BL/6J, NSG, B6(Cg)-Tyrc-2J/J, FVB/NJ, PWK/PhJ, 129S1/SvImJ, A/J, NOD/ShiLtJ, NZO/HILtJ, CAST/EiJ, and DBA/2J for rapid transgene insertion. Combined, this system enables predictable, rapid creation of precisely targeted transgenic animals across multiple genetic backgrounds, simplifying characterization, speeding expansion and use.

## Introduction

The mouse is a powerful and versatile research tool with which to gain a greater understanding of the human condition. One key capability is the ease and speed with which mice can be genetically modified, fueling the development of preclinical mouse models for drug development, human disease modeling and synthetic biology (1-5). However, it still remains technically challenging to integrate DNA constructs into the mouse genome *in vivo* in a precise and defined manner, which is essential for appropriate transgene control and expression (6). Although targeting nucleases, e.g., Zinc Finger Nucleases (ZFNs), Transcription Activator-Like Effector nucleases (TALENs), and CRISPR/Cas9 can mediate integration of exogenous DNA into the mouse genome in the zygote via homology-directed repair (HDR), the success of this approach is fraught with poorly defined variables and an inherent low efficiency of the intended event occurring, especially when the exogenous DNA exceeds a few kilobases (6-8).

Often where larger constructs are required to be added, researchers resort to the deceptively expedient solution of random (integration) transgenesis (9). Although this approach is well established and rapid, the resulting transgene insertions are haphazard and inefficient, often leading to unintentional genetic damage with potentially muddling downstream outcomes. For example, when donor constructs integrate into the genome, especially if prokaryotic donor sequences are retained, unpredicted local effects on host gene expression and silencing can occur (10, 11). Random transgenesis is also a misnomer. It has been shown that ∼50% of “random” transgene insertions disrupt coding sequences of endogenous genes, often invoking large deletions and structural variations at the integration regions (12, 13). Further, random transgenes frequently integrate as multiple copies, including concatemers, often leading to aberrant gene expression and transgene silencing (11, 14-17). Although such serendipitous heterogeneity of the resulting transgene expression can be of use, a significant amount of time is required to fully characterize the new strains, including multiple back-crosses to allow allelic segregation and to fully stabilize the phenotype.

Numerous approaches have been proposed to address this technology gap in genetic engineering, including lentiviral vectors and adeno-associated viruses (AAVs), as well as DNA transposons (e.g., Sleeping Beauty and piggyBac). For example, lentiviral-based systems can provide large payload capacity (18 kb) (18). However, they are associated with serious adverse effects (19), including genotoxicity (20) and immunogenicity (21). AAVs, on the other hand, have fewer adverse effects, but their payload capacity is limited to <5 kb (22). DNA transposases, such as Tn7 combined with catalytically inactive Cas9 for RNA-guided site-specificity, have been developed recently to perform targeted transgenesis of up to 10 kb of DNA, it is limited to prokaryotes (23-26). Further, transposases exhibit high off-target integration (27) requiring extensive engineering to achieve precise transgenesis in human cells, mice, and other animals. Lastly, severe DNA toxicity has also been observed (28, 29).

To overcome these limitations, we developed a precision transgenesis platform for integration of DNA constructs with minimal extraneous sequences directly using mouse zygotes. To minimize potential damage to the mouse genome our strategy uses the well-characterized safe harbor locus, Gt(ROSA)26Sor (ROSA26) with the pre-positioning of an integrase “landing pad” (attachment sites) for subsequent recombinant transgene integration. ROSA26 encodes a long non-coding RNA (lncRNA) under the control of a constitutive promoter. The region is transcriptionally active, with an open chromatin configuration leading to expression in all cell types examined, Further, its disruption does not lead to a reduction in fertility or other notable phenotypes (30-34). To facilitate the integration of large transgenes into ROSA26 we utilized the highly efficient serine site-specific recombinase (SSR) derived from the bacteriophage Bxb1 (35-37).

The Bxb1 mycobacteriophage was first isolated from *Mycobacterium smegmatis* in 1990, by Dr. William R. Jacobs, Jr., (see https://phagesdb.org/phages/Bxb1/) (36). Subsequently, it was determined that Bxb1 integrase (Int) is exquisitely suited to integrate DNA donor constructs into mammalian cells (38-41). Bxb1 Int-mediated recombination uses a minimal attachment site of 48 bp attP (attachment *Phage*) and a cognate 38 bp site attB (attachment *Bacterial*) between the donor and recipient DNAs. The attachment sites B and P represent four half-sites. After Bxb1 recombination, a half-site from each remains, yielding two 43bp sites (attL and attR, *Left* and *Right*) flanking the inserted DNA. Bxb1 Int requires no other secondary factors and provides irreversible unidirectional integration of donor DNAs under these conditions (42-44). Importantly, pseudo or intrinsic Bxb1 attachment sites that could lead to background off-target integrations (OTI) of donor DNA, have not been detected in the mouse genome (38, 41, 45-47). Furthermore, when fifteen integrases were compared in cell lines, Bxb1 Int was found to yield approximately two-fold greater and accurate recombinants when compared to the most widely studied recombinase φC31 (38, 46, 48).

Here we demonstrate that Bxb1 Int can efficiently and precisely introduce donor DNA constructs into the ROSA26 locus as single-copy transgenes in the desired orientation, directly in zygotes. In the first phase of this work, we used a *single* attachment site (attP-GT) in the mouse lines C57BL/6J (B6), FVB/NJ (FVB), and NOD.Cg-*Prkdc*^*scid*^ *Il2rg*^*tm1Wjl*^/SzJ (NSG), successfully generating recombinase mediated knock-in (RMKI) alleles using DNA minicircle donors to avoid integration of the prokaryotic backbone. In a second phase, we added a *second* heterologous attachment site (attP-GA) to the ROSA26 landing pad and show that this improved transgene insertion efficiency by more than 3-fold. Also, using dual sites enables larger donor DNAs to be inserted via recombinase mediated cassette exchange (RMCE), excluding the vector backbone and eliminating any need for a minicircle conversion step (see outline of approach in Fig. 1).

**Fig 1:**
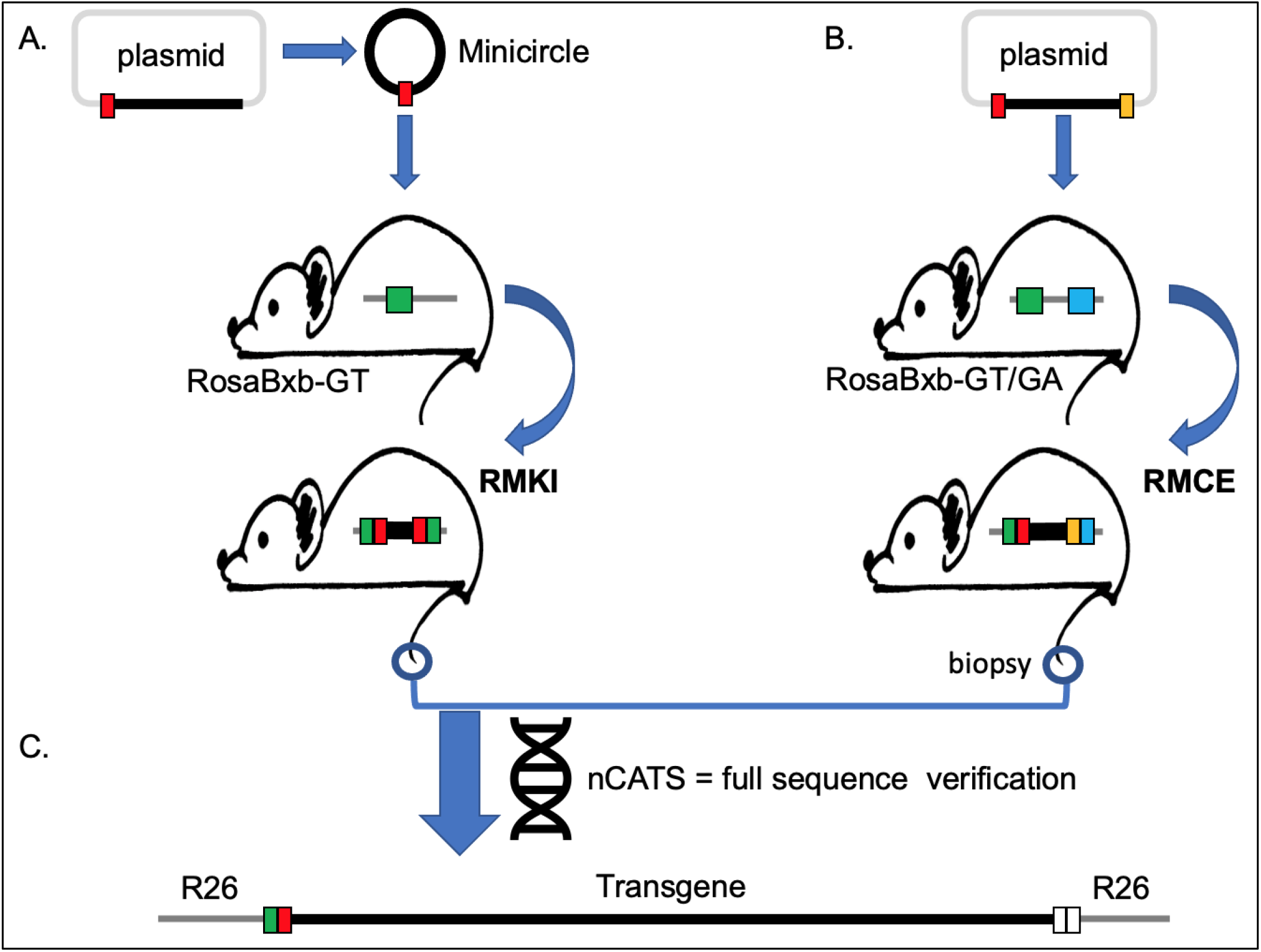
Overview of the Bxb1 Int-mediated transgenesis strategy. (A) Recombinase-Mediated Knock-In allele (RMKI) generated by in vivo recombination between vector free minicircle donor DNA carrying a single attB site and host mouse containing a single cognate attP site in the ROSA26 locus (RosaBxb-GT). (B) Recombinase-Mediated Cassette Exchange (RMCE) allele generated by in vivo recombination between plasmid DNA, carrying two heterologous attB sites and host mouse containing dual cognate attP sites (RosaBxb-GT/GA) in the ROSA26 locus. (C) Nanopore Cas9-targeted sequencing (nCATS) enables complete verification at the sequence level of the entire transgene including the adjacent genomic DNA (R26).

The Bxb1 Int-mediated transgenesis approach also provides distinct advantages for rapid screening and verification of the precise allele in its genomic context, enabling rapid and direct generation of *precision* transgenic animals in multiple genetic backgrounds. As proof of principle and to develop an important model for SARS-Cov2 research, we show the simultaneous creation of an identical Krt18-ACE2 transgenic allele on both B6 and NSG backgrounds, from construct ready to heterozygous N1 colonies in ∼4 months. We also present data from over 30 independent projects, providing full details of the Bxb1 Int-mediated transgenesis strategy and demonstrating the utility of this system for efficiently incorporating DNA construct from ∼3 kb to ∼43 kb into the mouse ROSA26 locus.

Finally, combining this strategy with nanopore Cas9-targeted sequencing (nCATS) enables us to completely sequence-verify the resulting transgenic allele rapidly *in its genomic context*. This complete strategy lends speed and confidence to the creation and characterization of genetically modified animal production with a level of precision and speed not previously obtainable.

## Materials and Methods

### Generation of RosaBxb-GT and RosaBxb-GT/GA Landing Pad alleles in host strains

Bxb1 attachment sites (attP) were inserted into the ROSA26 locus by CRISPR/Cas9 using TruGuide gRNAs (49) with donor oligos directly into zygotes. All donor oligonucleotides, CRISPR gRNA targeting sequences and PCR screening primers are listed in **Supplemental Table 1**. Microinjection was performed as described previously by (50), or by electroporation (51, 52). Where individual experimental conditions varied, these are detailed in Table 1.

**Table 1:**
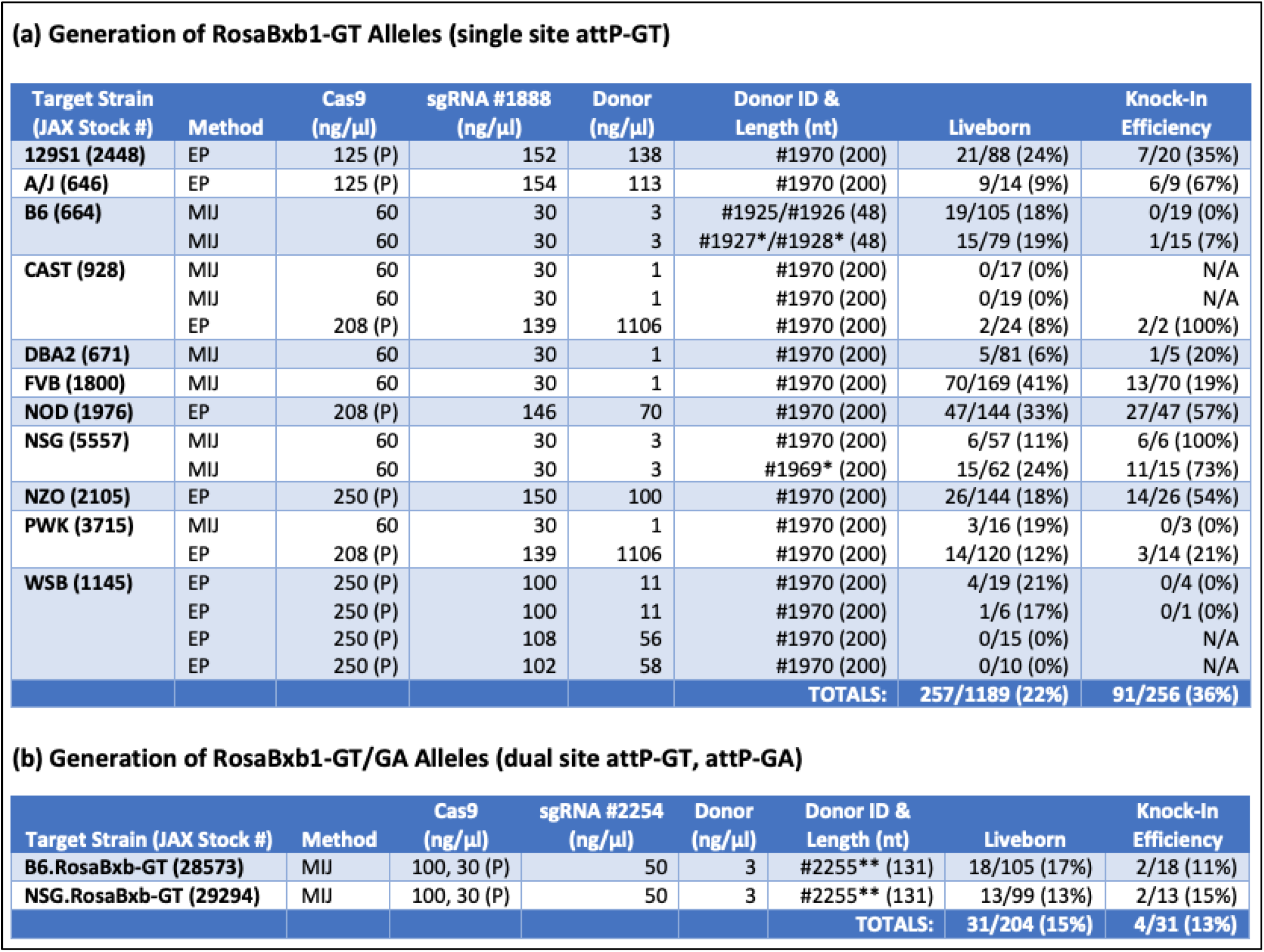
Summary of CRISPR/Cas9-mediated Knock-In efficiency of Bxb1 attP Site(s) into the ROSA26 locus. shows the range of CRISPR/Cas9 conditions used to generate the RosaBxb-GT and RosaBxb-GT/GA alleles on multiple genetic backgrounds. MIJ, microinjection; EP, electroporation are listed and all oligonucleotides were phosphothioated unless noted (*). Cas9 was delivered as mRNA, except where indicated as Protein, (P). All 200nt oligo donors contained the attP site at the center, i.e., 76nt of homology to the ROSA26 locus on either side. Liveborn is the number of animals available to screen at wean over the number of embryos transferred following EP/MIJ. KI efficiency refers to the correct integration of the attP site into the ROSA26 locus. For B6 *only* we used a GUIDE-seq strategy with a dsDNA donor. The attP-GT was inserted into the ROSA26 locus into the same place and orientation in all genetic backgrounds using Cas9 guide #1888. The attP-GA site was inserted 240bp 3’ of the attP-GT site in previously-modified B6.RosaBxb-GT and NSG.RosaBxb-GT strains with guide #2255.

(a) Generation of RosaBxb1-GT Alleles (single site attP-GT)
| Target Strain (JAX Stock #) | Method | Cas9 (ng/μl) | sgRNA #1888 (ng/μl) | Donor (ng/μl) | Donor ID & Length (nt) | Liveborn | Knock-In Efficiency |
| --- | --- | --- | --- | --- | --- | --- | --- |
| 129S1 (2448) | EP | 125 (P) | 152 | 138 | #1970 (200) | 21/88 (24%) | 7/20 (35%) |
| A/J (646) | EP | 125 (P) | 154 | 113 | #1970 (200) | 9/14 (9%) | 6/9 (67%) |
| B6 (664) | MIJ | 60 | 30 | 3 | #1925/#1926 (48) | 19/105 (18%) | 0/19 (0%) |
|  | MIJ | 60 | 30 | 3 | #1927*/#1928* (48) | 15/79 (19%) | 1/15 (7%) |
| CAST (928) | MIJ | 60 | 30 | 1 | #1970 (200) | 0/17 (0%) | N/A |
|  | MIJ | 60 | 30 | 1 | #1970 (200) | 0/19 (0%) | N/A |
|  | EP | 208 (P) | 139 | 1106 | #1970 (200) | 2/24 (8%) | 2/2 (100%) |
| DBA2 (671) | MIJ | 60 | 30 | 1 | #1970 (200) | 5/81 (6%) | 1/5 (20%) |
| FVB (1800) | MIJ | 60 | 30 | 1 | #1970 (200) | 70/169 (41%) | 13/70 (19%) |
| NOD (1976) | EP | 208 (P) | 146 | 70 | #1970 (200) | 47/144 (33%) | 27/47 (57%) |
| NSG (5557) | MIJ | 60 | 30 | 3 | #1970 (200) | 6/57 (11%) | 6/6 (100%) |
|  | MIJ | 60 | 30 | 3 | #1969* (200) | 15/62 (24%) | 11/15 (73%) |
| NZO (2105) | EP | 250 (P) | 150 | 100 | #1970 (200) | 26/144 (18%) | 14/26 (54%) |
| PWK (3715) | MIJ | 60 | 30 | 1 | #1970 (200) | 3/16 (19%) | 0/3 (0%) |
|  | EP | 208 (P) | 139 | 1106 | #1970 (200) | 14/120 (12%) | 3/14 (21%) |
| WSB (1145) | EP | 250 (P) | 100 | 11 | #1970 (200) | 4/19 (21%) | 0/4 (0%) |
|  | EP | 250 (P) | 100 | 11 | #1970 (200) | 1/6 (17%) | 0/1 (0%) |
|  | EP | 250 (P) | 108 | 56 | #1970 (200) | 0/15 (0%) | N/A |
|  | EP | 250 (P) | 102 | 58 | #1970 (200) | 0/10 (0%) | N/A |
| TOTALS: |  |  |  |  |  | 257/1189 (22%) | 91/256 (36%) |

**Table 1: Summary of CRISPR/Cas9-mediated Knock-In efficiency of Bxb1 attP Site(s) into the ROSA26 locus** shows the range of CRISPR/Cas9 conditions used to generate the RosaBxb-GT and RosaBxb-GT/GA alleles on multiple genetic backgrounds.
| Target Strain (JAX Stock #) | Method | Cas9 (ng/μl) | sgRNA #2254 (ng/μl) | Donor (ng/μl) | Donor ID & Length (nt) | Liveborn | Knock-In Efficiency |
| --- | --- | --- | --- | --- | --- | --- | --- |
| B6.RosaBxb-GT (28573) | MIJ | 100, 30 (P) | 50 | 3 | #2255** (131) | 18/105 (17%) | 2/18 (11%) |
| NSG.RosaBxb-GT (29294) | MIJ | 100, 30 (P) | 50 | 3 | #2255** (131) | 13/99 (13%) | 2/13 (15%) |
| TOTALS: |  |  |  |  |  | 31/204 (15%) | 4/31 (13%) |

For all strains created here, insertion of the initial Bxb1 attP-GT site used CRISPR/Cas9 with gRNA #1888, targeting the ROSA26 locus. For only mouse strain B6, a CRISPR/Cas9 GUIDE-seq strategy using duplexed donor oligos #1927/#1928 was applied (53). For all other strains insertion of the initial Bxb1 attP-GT site used CRISPR/Cas9 mediated HDR with single-stranded donor oligo #1969 or #1970. The second Bxb1 site attP-GA, was inserted into zygotes isolated from B6 or NSG RosaBxb-GT mice homozygous for the initial Bxb1 attP-GT site and used CRISPR/Cas9 mediated HDR with gRNA #2254 and donor oligo #2255. The resulting two 48bp attP sites are positioned 240bp apart, with the initial attP-GT site placed 4bp from the *XbaI* site in intron 1 of the ROSA26 locus. In all cases following microinjection or electroporation zygotes were transferred to pseudopregnant females and brought to term.

Tissue from tail tip or ear punch was used to isolate DNA for PCR analysis as outlined in (50). Specifically for the first attP-GT site we used PCR primers #1699 and #2162, yielding a 254bp product if the RosaBxb-GT allele was present. For the detection of the second site attP-GA, PCR using primers #2161 and #2162 yielded a 336bp product if the RosaBxb-GT/GA allele was present. To fully validate the modified ROSA26 locus, PCR using primers #1699 & #1703 flanking the entire modified region were used followed by Sanger sequencing. After one back-cross to the unmodified parental strain, lines confirmed to be carrying the attP site/s were inbred to establish homozygous colonies as listed in the Results. The RosaBxb-GT/GA alleles on B6-albino and NOD.ShiLtJ backgrounds were produced by selective back-crossing of B6.RosaBxb-GT/GA mice with B6(Cg)-*Tyr*^*c-2J*^/J (stock#58), and NSG.RosaBxb-GT/GA with NOD/ShiLtJ (stock#1976), respectively.

### RMKI host donor plasmid construction

To prepare minicircle DNA, desired transgenes are first cloned into a donor host plasmid (p5087). This plasmid was derived from the MN530A-1 (System Biosciences) by replacing the GFP Reporter sequence with a multiple cloning site using the restriction enzymes *XmaI* and *StuI* and duplexed donor oligonucleotides #2020 and #2021 (**Supplemental Table 1**). This plasmid was further modified to include the Bxb1 attB-GT site using restriction enzymes *XbaI* and *EcoRI* and duplexed oligonucleotides #2025 and #2026. Donor plasmid DNA (containing the attB-GT and the desired transgene) is prepared for injection by conversion into a vector-free minicircle using the MC-Easy™ Minicircle DNA Production Kit (SystemBio). Minicircle DNA is then isolated using the Purelink HiPure Plasmid Filter Midiprep kit (Invitrogen). To ensure complete removal of the parental plasmid, a restriction digest is performed to exclusively linearize the parental plasmid followed by an overnight digestion with Plasmid-Safe™ ATP-Dependent DNase (Lucigen). Prior to MIJ, the resultant minicircle DNA is *extensively* purified by phenol-chloroform extraction, precipitated with ethanol-sodium acetate and reconstituted in nuclease-free 10mM Tris/0.1mM EDTA/pH7.5.

### RMKI by Bxb1 Int using RosaBxb-GT Host Mice

Recombinase-Mediated Knock-In (RMKI) of donor DNA into the single-site RosaBxb-GT host strains is achieved by microinjection of 1-30ng/µl donor DNA with 100ng/µl Bxb1 mRNA (Trilink) into zygotes of the desired attP-GT carrier strains. Typically, host zygotes heterozygous for Bxb1-attP are used, as these can be readily generated by super-ovulating wild-type dams, using for example B6 mated to homozygous studs, B6.RosaBxb-GT.

### RMCE donor plasmid construction

For RMCE alleles, DNA vector requires the Bxb1 attB-GT and Bxb1 attB-GA sites are positioned in the correct orientation and flanking the desired sequence. Our donor plasmid p5154 (Addgene reference 175390) is intended for small-medium sized donor DNAs (<15kb) while the p5155 (Addgene reference 175391) is a low-copy version for use with larger donor DNAs (>15kb). Prior to MIJ, plasmid DNA containing the requisite attB-GT and attB-GA sites flanking the transgenic donor DNAs are isolated using the Purelink HiPure Plasmid Filter Midiprep kit (Invitrogen) followed by *extensive* purification via phenol-chloroform extraction, ethanol-sodium acetate precipitation with a final reconstitution in nuclease-free 10mM Tris/0.1mM EDTA/pH7.5.

### RMCE in RosaBxb-GT/GA Host Mice

Recombinase-Mediated Cassette Exchange (RMCE) of donor DNA into the dual-site RosaBxb-GT/GA host strains is achieved by microinjection of 1-30ng/µl donor DNA with 100ng/µl Bxb1 mRNA (Trilink) into zygotes of the desired strain. Generally, host zygotes heterozygous for the dual Bxb1-attP sites were generated by super-ovulating wild-type dams, e.g., B6 mated to B6.RosaBxb-GT/GA homozygous studs. Microinjection preps were prepared in nuclease-free 10mM Tris/0.1mM EDTA/pH7.5 (IDT).

### Screening offspring for RMCE and RMKI donor integration

A generalized reductive sequential screening strategy for identifying all Bxb1 Int-mediated alleles was developed using a series of four to six simple PCR assays on crude DNA lysates prepared from tail-tip or ear punch biopsies (50). As outlined in Fig. 2, these assays are designed to rapidly identify the correctly integrated alleles as well as screen for founder animals that also carry random insertions of donor DNA. The PCR primers used are defined in **Supplemental Table 1**. It is strongly recommended that time be taken to optimize these screening primers, see (50).

**PCR1 Assay** is designed to rapidly identify candidate Founders using a generic multiplex PCR with primers #1699, #1703 and #2164. PCR1 also verifies the integrity of the genomic DNA template, yielding a wild type band of 776bp, simultaneously generating a unique band in samples where the recombination event between attP-GT and attB-GT has occurred (∼252bp); see Fig. 2. This PCR is performed on all potential Founder animals.
**PCR2 Assay** is a transgene-specific PCR using gene sequence-specific primers (GSP) designed to target a unique portion of the donor and is performed on all candidate offspring. Note, while this assay does not differentiate between random transgene events and correctly integrated alleles, its increased sensitivity will detect candidates that the multiplex PCR1 may miss due to low copy number.
**PCR3 Assay**: Founder animals that show a positive result by PCR1 or PCR2 are next screened using two project-specific In/Out PCR assays designed to amplify the allele across both left and right ROSA26/Transgene junctions. As shown in Fig. 2, the In/Out-Left (IOL) PCR uses a forward primer in ROSA26 5’ of the attP-GT site (e.g., primer #1699) with a transgene-specific reverse primer (GSP) that is 3’ of the attB-GT site. Similarly, the In/Out-Right (IOR) PCR assay uses a reverse primer in ROSA26 3’ of the attP site (e.g., primer #1703) with a matching donor-specific forward primer. Resulting PCR products are sequenced to confirm the correct junction between host and donor DNA resulted (50). Where possible (∼<10kb), these assays are modified to enable full-sequence verification using long-range PCR. In these instances, the In/Out PCRs are re-designed so as to overlap, i.e., the forward primer for the IOR PCR should be 5’ of the reverse primer for the IOL PCR.
**PCR4 Assay**: To identify random transgenesis in putative candidate Founders, project-specific PCR for Off-Target Integration (OTI) are performed. These assays are designed to target across the attB site(s) in the *donor* DNA using vector backbone-specific primers (BB) with gene matching transgene-specific primers (GSP). Products from one or both of these assays indicate either a random insertion or an aberrant integration at the landing pad site has occurred.

**Fig 2:**
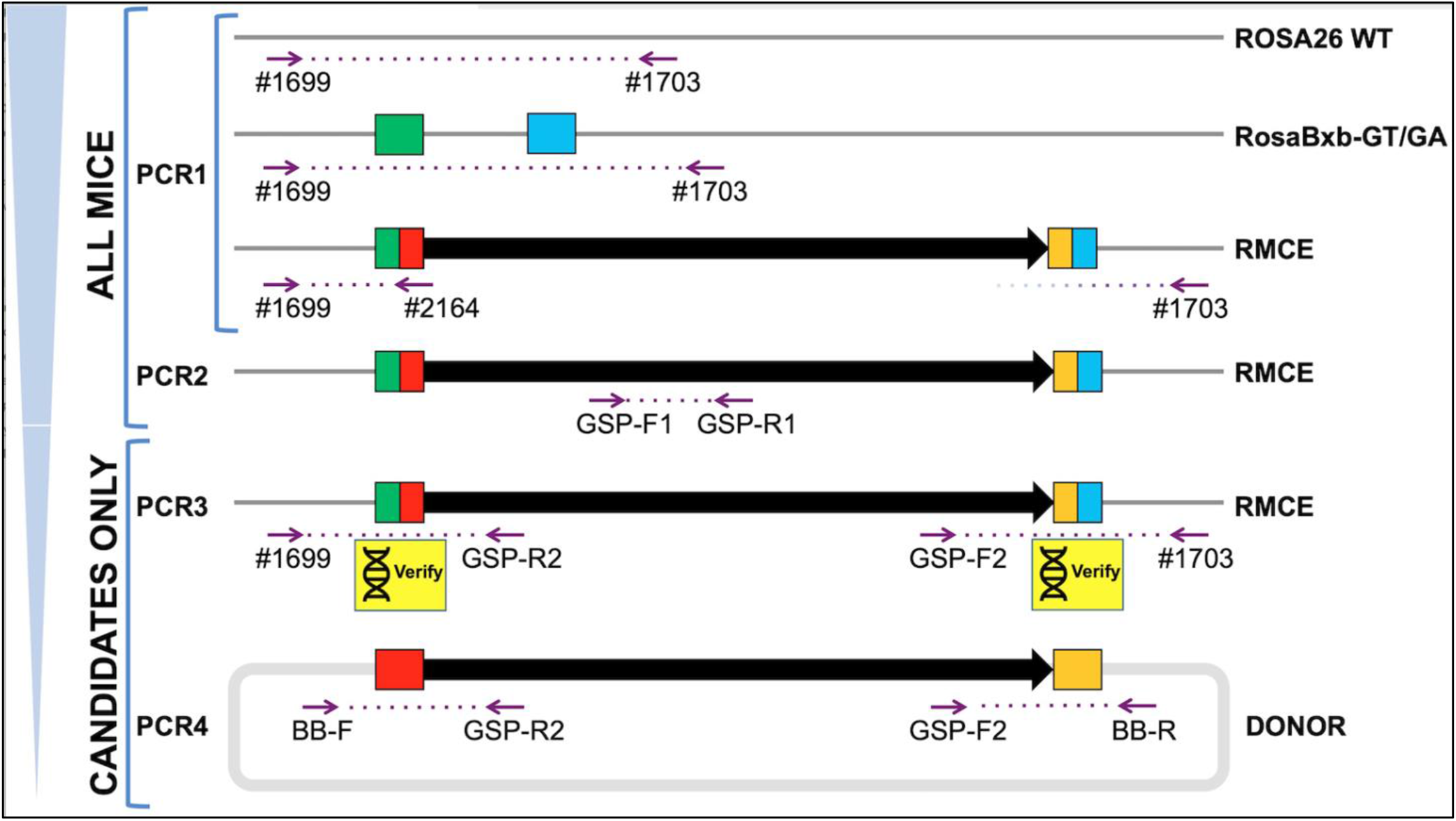
RMCE Screening Strategy. Screening offspring generated by microinjection follows a reductive regime identifying candidate animals. Both PCR1 and PCR2 assays are used to screen *all* potential Founder animals, while PCR3 and PCR4 assays are performed on only those positive by PCR1 and/or PCR2. A “generic” multiplex PCR1 is used to identify the presence of the RMCE allele resulting from *any* DNA donor, targeting both the ROSA26 locus and the recombined attR-GT site (green/red box). PCR1 serves as a DNA template quality control and is later used to determine zygosity when verified animals are inbred. The project-specific PCR2, using transgene-specific forward and reverse primers (GSP), verifies that donor DNA is present in the sample (it is not designed to distinguish between random insertion events and the RMCE allele). PCR3 assays verify that the correct (ROSA26) integration event has occurred utilizing an In/Out strategy. These PCRs span each newly created unique junction sites and are followed by Sanger Sequencing verification. For transgenes <10Kb, re-positioning of GSP-R2 and GSP-F2 can be used to generate overlapping PCR products that allow sequence verification of the entire allele from the two long-range amplicons. Assay PCR4 is performed to screen for the presence of aberrant/random insertion using vector-backbone specific primers (BB) combined with transgene specific primers (GSP) designed to span the attB sites in the DNA donor.

RMKI alleles generated by minicircle integration into a single attP-GT landing pad in ROSA26 are screened in the same manner as RMCE alleles, except only one PCR4 Assay is needed to span the single attB site in the minicircle donor. However, an additional PCR is recommended to screen for the presence of any parental plasmid backbone that may have eluded the minicircle production process. These assays are project-specific and should use primers spanning the φC31 and attB sites in the parental plasmid (not shown).

Following the RMKI/RMCE strategy outlined here, Founder animals are expected be heterozygous for any Bxb1 Int-mediated integration. However, donor DNA integration may not occur with 100% efficiency in the single cell zygote, leading to a mosaic founder animal whose actual zygosity will be less than 50%. To confirm germline transmission and establish the integrity of the allele, one to four carrier Founders are back-crossed to the wild type parental strain (*never* other Founders). The resulting N1 animals are then subjected to the same PCR screening process (including PCR4), regardless of the genotype of the mosaic Founder animal. We recommend that a single verified N1 animal then be used to establish the new strain. Consequently, subsequent genotyping requires only a single multiplex PCR (e.g., IOL PCR3 with primer #1703 included) to determine the presence and zygosity of the new allele.

### nCATS and data visualization

Unique gRNA’s for nanopore Cas9-targeted sequencing (nCATS) are designed 5’ and 3’ of the region to be sequenced (see **Supplemental Table 1**). For genomic DNA extraction, tissue is pulverized on dry ice in a Bessman Tissue Pulverizer (Spectrum™, 189476). High molecular weight gDNA extraction using a Monarch® HMW DNA Extraction Kit for Tissue (New England Biolabs, T3060L) is performed on tissues isolated from N1 or higher generation mice. We do not recommend this process on founder animals as they are both often mosaic and potentially heterogeneous for the modification. nCATS targeted libraries are constructed as detailed in (54) with the exception of ACE2-CDS of B6.Cg-Tg(K18-ACE2)2Prlmn/J JAX# 034860, where a single cut and read library was made per ACE2-CDS targeting gRNA, and pooled prior to Ampure bead purification. nCATS libraries are routinely sequenced for 24 hours on a GridION using a R9.4.1 flow cell (Oxford Nanopore Technologies). After 24 hours of sequencing, flow cells are nuclease flushed (Flow Cell Wash Kit EXP-WSH004, Oxford Nanopore Technologies) and reloaded with new nCATS libraries up to three times.

Base calling is done using GUPPY (v3.2.10) and the resulting FASTQ files are aligned to a reference sequence using minimap2(v2.17). Custom reference sequences were constructed for transgene insertion sites using the Mus musculus C57BL/6J reference genome (MM10) and corresponding insert sequence. Alignment results are subject to MapQ score filtering using Samtools (v1.11). Subsequent coverage depth for on-target reads are generated using Samtools (v1.11) and Bedtools (v2.29.2). On-target reads are visualized using the Integrative Genomics Viewer (IGV) and consensus sequences were generated using Medaka (v1.0.3) and annotated in Snapgene (v5.3).

## Results

The dual attP site landing pad alleles were constructed over two phases, enabling testing of multiple CRISPR/Cas9 based strategies for optimal knock-in allele generation. After establishing the single site attP allele on various backgrounds, the B6 and NSG versions were re-targeted to add a second attP site in *cis*. Both single and dual-site mouse strains were used to generate precision transgenics via Bxb1-mediated recombination. Three single site strains (B6, FVB, NSG) used donor DNA as minicircles to generate vector-free RMKI (recombinase-mediated knock-in) alleles. In contrast, the two (B6 and NSG) dual-site strains used plasmids or modified BACs to generate vector-free RMCE (recombinase-mediated cassette exchange) alleles. Compared to the single site, the dual-site strains significantly improve average efficiency from 5% to 15%, while also enabling insertion of larger donor DNA (to at least 43 kb).

### Generation of single site Bxb1 attP-GT landing pad mouse strains (RosaBxb-GT)

Initially, a single Bxb1 attP-GT site was introduced 3’ of the *XbaI* site in intron 1 of the ROSA26 locus (see Fig. 3). To reduce the potential for off-target cutting, we used an 18-mer TRU-guide gRNA (49) which consistently cuts at ≥85% at this locus. For the mouse strain B6 only, we used a GUIDE-seq strategy (53) introducing the attP site using Cas9 mRNA, gRNA#1888 and a 48bp dsDNA oligonucleotide donor (duplexed oligos #1927/#1928) delivered into B6 zygotes by microinjection. Analysis of the resulting allele in B6 showed successful integration of the complete attP-GT site into the ROSA26 locus. However, it was evident that this approach was inefficient (1/15) and in this instance led to a lesion at the integration site, with a +6bp/-3bp INDEL preceding the 5’ end of the attP-GT site; see Fig. 3. As a consequence, this strategy was abandoned and for all other attP integrations into other strains of mice strains used Cas9 mediated HDR with a single-stranded oligo (#1969 or #1970) which included homology arms (12). These data are outlined in **Table 1**.

**Fig 3:**
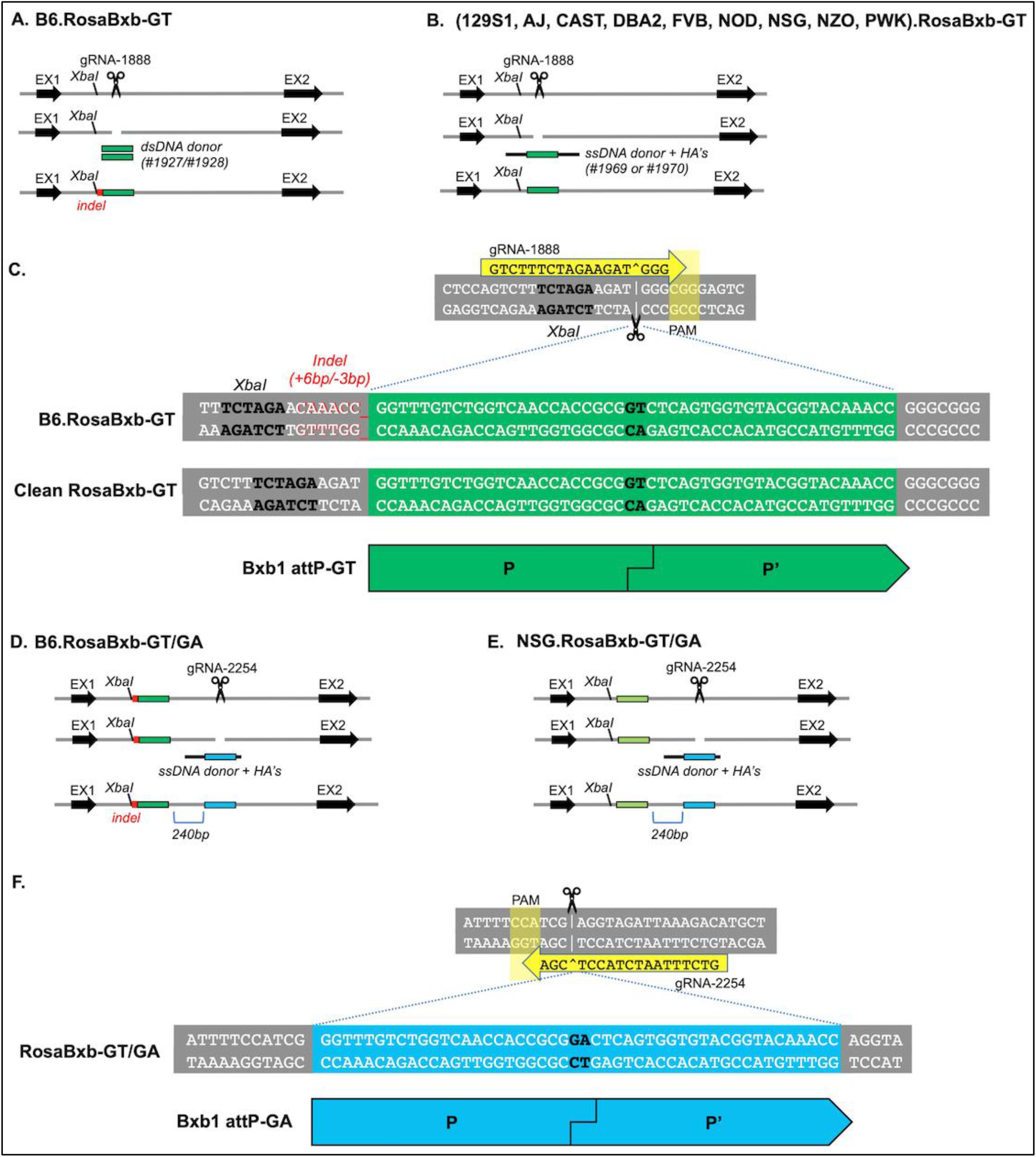
Construction of RosaBxb-GT alleles on multiple genetic backgrounds. (A) For the mouse strain B6 only, a CRISPR/Cas9 GUIDE-seq strategy using duplexed donor oligos #1927 and #1928 and guide gRNA #1888 resulted in the insertion of the attP-GT site with an indel. (B) A CRISPR/Cas9 HDR strategy with donor oligos #1969 or #1970 and gRNA #1888 was used for all other backgrounds, resulting in scarless incorporation of the attP site to generate the RosaBxb-GT alleles. (C) Sequence of Bxb1 attP-GT site for B6.RosaBxb1-GT, showing the indel that occurred in B6 and the correct RosaBxb-GT allele with perfect integration that resulted in mouse strains 129S1.RosaBxb-GT, AJ.RosaBxb-GT, CAST.RosaBxb-GT, DBA2.RosaBxb-GT, FVB.RosaBxb-GT, NOD.RosaBxb-GT, NSG.RosaBxb-GT, NZO.RosaBxb-GT, and PWK.RosaBxb-GT. Generation of RosaBxb-GT/GA alleles on B6 (D) and NSG (E) genetic backgrounds using a CRISPR/Cas9 HDR strategy wherein the attP-GA site was placed 240bp 3’ of the attP-GT site, using gRNA #2254 and asymmetric donor oligo #2255. (F) Close-up of the Bxb1 attP-GA site region in strains B6.RosaBxb1-GT/GA and NSG.RosaBxb1-GT/GA.

As shown in **Table 1**, all strain backgrounds attempted resulted in successful generation of the ROSA26 attP-GT allele, with the exception of WSB. The primary challenge for the CAST and WSB strains was low embryo production (due to super-ovulation resistance) combined with poor embryo survival post-manipulation. For CAST, this was overcome using timed matings and electroporation of zygotes to introduce duplexed Cas9 protein + guide and donor oligo, producing two carrier Founders. Unfortunately, while CRISPR/Cas9 targeted NHEJ was detected in WSB Founders, we did not detect knock-in of the attP site in the very limited number of offspring obtained. For all other strains listed, the attP knock-in allele was detected, transmitted and confirmed as described. After a minimum of one back-cross to the appropriate strain background, each new strain was bred to homozygosity. Ultimately a single line was selected for each genetic background and a list of all strains are shown in

### Use of single site RosaBxb-GT strains to generate RMKI alleles

The generation of vector-free RMKI alleles was achieved as shown in **Fig. 4**, with the screening strategy as outlined in **Fig. 2**. A range of conditions were attempted to optimize the system for efficient generation of RMKI alleles with three single-site strains, B6.RosaBxb-GT, FVB.RosaBxb-GT, and NSG.RosaBxb-GT. Results from all conditions tested for each donor minicircle were combined and summarized in **Table 3**. This table highlights our attempts to integrate twelve unique minicircle plasmid donors ranging in size from 3.2 to 9.9kb. The highly variable results were expected due to the multiple conditions tested. Overall, we demonstrated that more than half (22/39) of the individual

**Fig 4:**
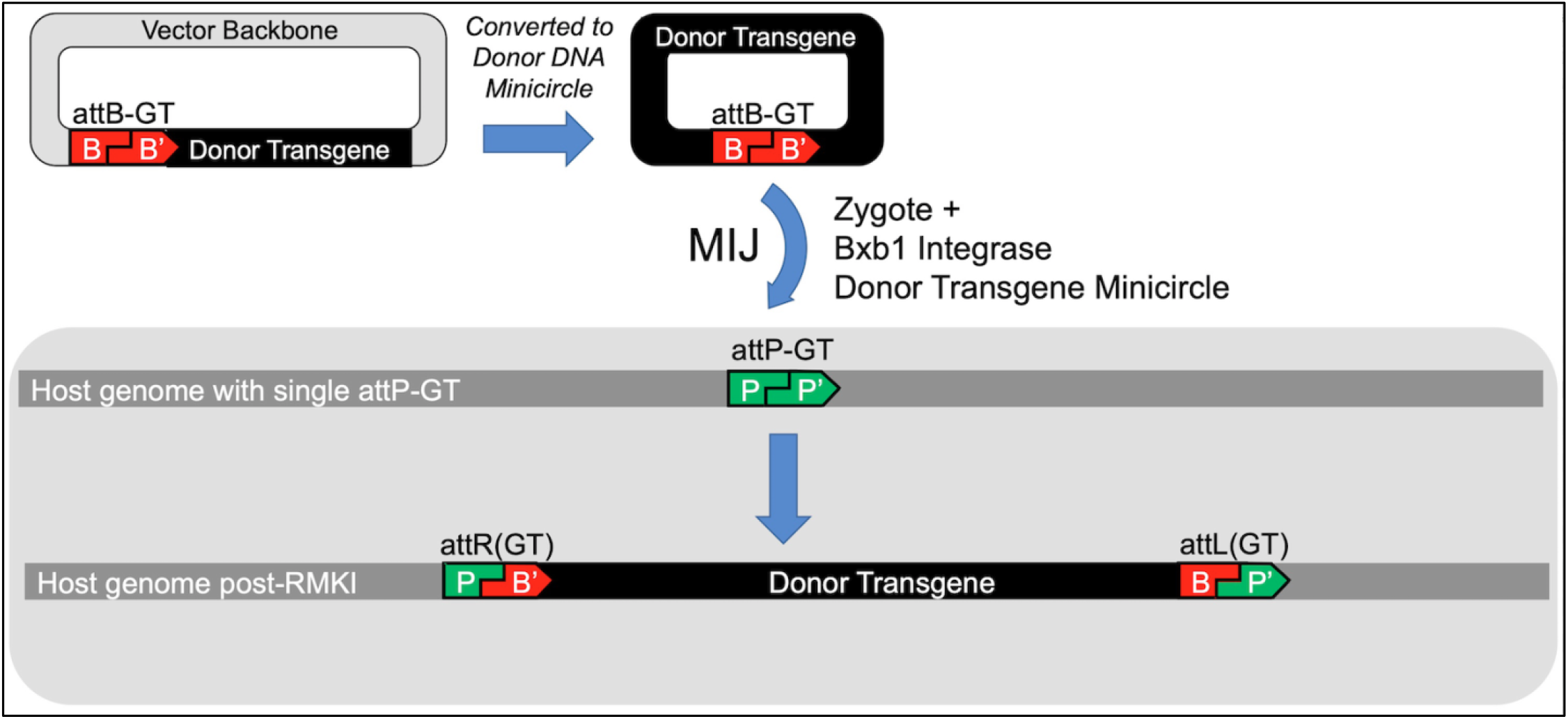
Schema for using single attP-GT site mouse strains targeted for recombinase mediated knock-in transgenesis (RMKI). Before microinjection, the plasmid vector containing the donor transgene and a single cognate attB-GT site are converted into (vector-less) DNA minicircles and purified. Along with Bxb1 Int, the minicircles are delivered into zygotes carrying the attP-GT landing pad allele, enabling precision transgene integration as the attP (host genome), and attB (donor DNA) sites recombine to form attR and attL sites.

In <5% of identified integrations, we detected aberrant integration events when analyzing RMKI alleles. These appear to be the result of unintended rearrangement occurring during donor DNA minicircle production. Analysis of these alleles suggests that *in vitro* fC31-based recombination designed to aid minicircle production has combined multiple plasmid DNAs producing some minicircles with 2-3 copies in tandem of the intended transgene. Where identified, these aberrant alleles are shown in **Table 3**. In the case of MC-1, which resulted in 3 copies of a double-inverted open-reading frame (DIO) allele designed to express a toxin only in cells where Cre is expressed, the allele was maintained and shown to function as intended (data not shown). As a general rule however, such aberrant allele lines were detected and discarded early in the screening process, or after sequencing the region by nCATS (data not shown).

**Table 2:** Summary of strains developed and available from JAX. A list of all mouse strains described in this paper and available from the Jackson Laboratory, showing their short nickname, JAX Stock Number, and the original genetic background for each new line. Note: mouse lines B6A.RosaBxb-GT/GA and NOD.RosaBxb-GT/GA (JAX stock number 036153 and 036181) were established by backcross and allelic segregation, while all other attP landing pad strains were created by MIJ or EP of the source strain indicated. The two Krt18-ACE2 alleles (JAX stock numbers 036363 and 035893) were created by RMCE, resulting from microinjection of the same donor DNA into the B6 and NSG dual attP strains (036152 and 036151).

| Single Site Landing Pad Strains (RosaBxb-GT) for RMKI |  |  |  |
| --- | --- | --- | --- |
| Source Strain (Stock #) | JAX Stock # | Full Nomenclature | Strain Nickname |
| 129S1/SvImJ (2448) | 35029 | 129S1/SvImJ-Gt(ROSA)26Sor<sup>em12Mvw</sup>/MvwJ | 129S.RosaBxb-GT |
| A/J (646) | 35275 | A/J-Gt(ROSA)26Sor<sup>em18Mvw</sup>/MvwJ | AJ.RosaBxb-GT |
| C57BL/6J (664) | 28573 | C57BL/6J-Gt(ROSA)26Sor<sup>em2Mvw</sup>/MvwJ | B6.RosaBxb-GT |
| CAST/EiJ (928) | 35033 | CAST/EiJ-Gt(ROSA)26Sor<sup>em16Mvw</sup>/MvwJ | CAST.RosaBxb-GT |
| DBA/2J (671) | 35031 | DBA/2J-Gt(ROSA)26Sor<sup>em14Mvw</sup>/MvwJ | DBA2.RosaBxb-GT |
| FVB/NJ (1800) | 35396 | FVB/NJ-Gt(ROSA)26Sor<sup>em20Mvw</sup>/MvwJ | FVB.RosaBxb-GT |
| NOD/ShiLtJ (1976) | 35320 | NOD/ShiLtJ-Gt(ROSA)26Sor<sup>em11Mvw</sup>/MvwJ | NOD.RosaBxb-GT |
| NSG (5557) | 29294 | NOD.Cg-Gt(ROSA)26Sor<sup>em3Mvw</sup> Prkdc<sup>scid</sup> Il2rg<sup>tm1Wjl</sup>/MvwJ | NSG.RosaBxb-GT |
| NZO/HILtJ (2105) | 36178 | NZO/HILtJ-Gt(ROSA)26Sor<sup>em31Mvw</sup>/MvwJ | NZO.RosaBxb-GT |
| PWK/PhJ (3715) | 35276 | PWK/PhJ-Gt(ROSA)26Sor<sup>em19Mvw</sup>/MvwJ | PWK.RosaBxb-GT |

| Dual Site Landing Pad Strains (RosaBxb-GT/GA) for RMCE |  |  |  |
| --- | --- | --- | --- |
| Source Strain/s (Stock #) | JAX Stock # | Full Nomenclature | Strain Nickname |
| B6.RosaBxb-GT (28573) | 36152 | C57BL/6J-Gt(ROSA)26Sor<sup>em7Mvw</sup>/MvwJ | B6.RosaBxb-GT/GA |
| B6(Cg)-Tyrc-2J/J (58),<br>B6.RosaBxb-GT (28573) | 36153 | B6.Cg-Gt(ROSA)26Sor<sup>em7Mvw</sup> Tyr<sup>c-2J</sup>/MvwJ | B6A.RosaBxb-GT/GA |
| NSG.RosaBxb-GT (29294) | 36151 | NOD.Cg-Gt(ROSA)26Sor<sup>em5Mvw</sup> Prkdc<sup>scid</sup> Il2rg<sup>tm1Wjl</sup>/MvwJ | NSG.RosaBxb-GT/GA |
| NOD/ShiLtJ (1976),<br>NSG.RosaBxb-GT (29294) | 36181 | NOD(Cg)-Gt(ROSA)26Sor<sup>em5Mvw</sup>/MvwJ | NOD.RosaBxb-GT/GA |

**Table 2: Summary of strains developed and available from JAX**
| Available Mouse Strains Generated by RMCE |  |  |  |
| --- | --- | --- | --- |
| Source Strain/s (Stock #) | JAX Stock # | Full Nomenclature | Strain Nickname |
| B6.RosaBxb-GT/GA (36152) | 36363 | C57BL/6J-Gt(ROSA)<sup>26Sorem25(KRT18-ACE2)</sup>Mvw/MvwNrosJ | B6.RMCE[Krt18-ACE2] |
| NSG.RosaBxb-GT/GA (36151) | 35893 | NOD.Cg-Gt(ROSA)<sup>26Sorem27(KRT18-ACE2)</sup>Mvw Prkdc<sup>scid</sup> Il2rg<sup>tm1Wjl</sup>/MvwJ | NSG.RMCE[Krt18-ACE2] |

**TABLE 3.**
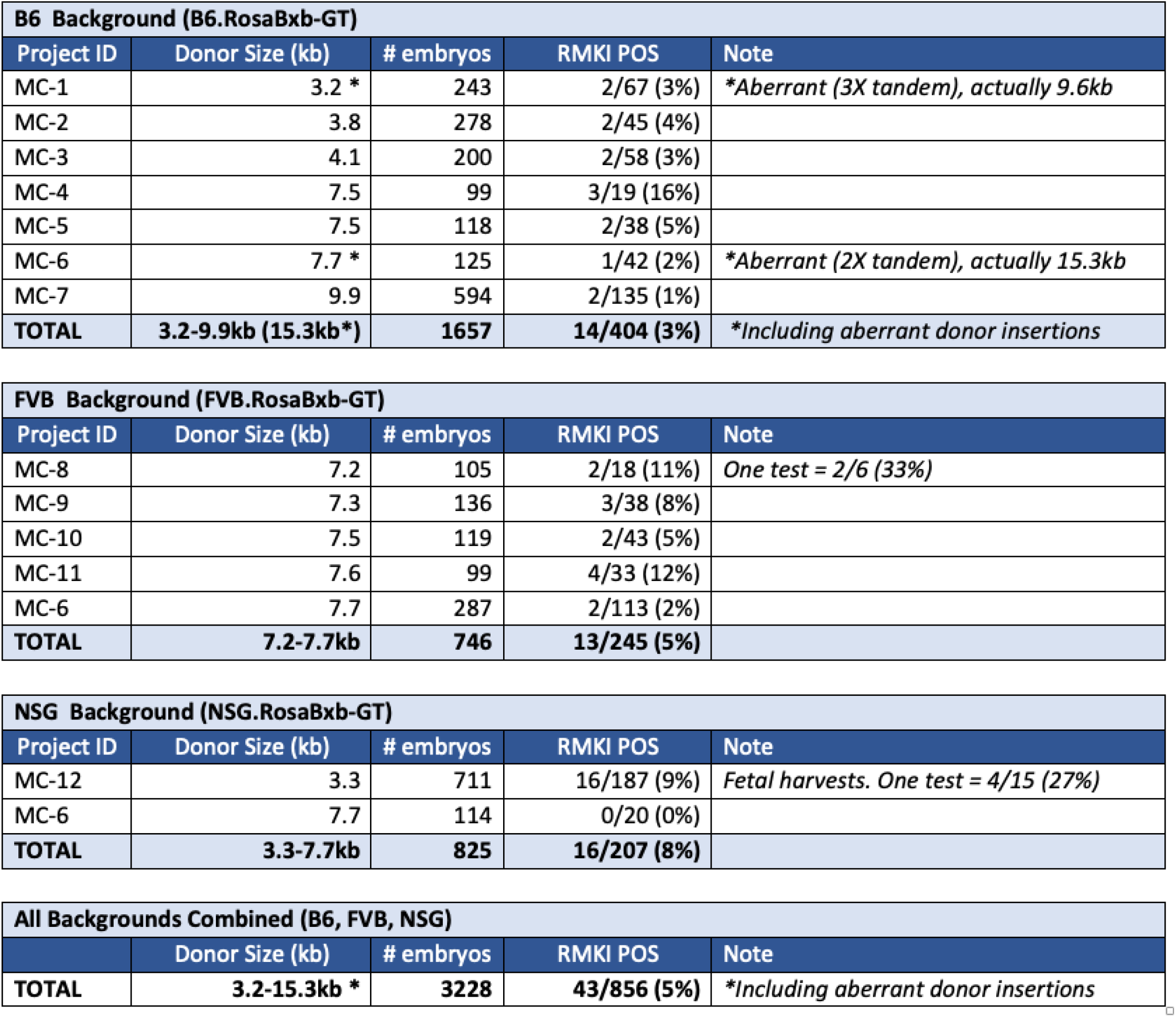
Summary of RMKI attempts. Summary of fourteen RMKI projects attempting to integrate twelve unique minicircles into the RosaBxb-GT single site allele across three different genetic backgrounds (B6, FVB, NSG). Minicircle DNA donor size, the number of microinjected embryos transferred and the number of positive Founders over the number of animals screened is shown. Recommended conditions are given in Material and Methods. Note, results from multiple conditions using the same donor were combined. Individual conditions tested ranged from 0-33% correct integrations. Aberrant alleles resulting from the minicircle production step occasionally resulted in larger inserts than intended (indicated by *). All projects taken to liveborn and established colonies, except for MC-12, which was used solely to optimize conditions and were collected as embryos for rapid analysis.

### Addition of second attP-GA site

Although the RMKI strategy was successful with inserts up to ∼15kb, the overall efficiency of donor construct integration, plus operational constraints in making minicircle donor DNA led us to engineer RMCE enabled host mice. This was achieved through the addition of a second (heterologous) attP site, with “GA” as the central dinucleotide as opposed to the wild type “GT” used for the first site. This second site (attP-GA) was placed in *cis*, 240bp 3’ of the first attP-GT site, in B6.RosaBxb-GT and NSG.RosaBxb-GT strains (**see Fig. 3**). This sequential modification was achieved using CRISPR/Cas9 and sgRNA#2254 to mediate the HDR integration of donor oligo #2255 into zygotes homozygous for the initial attP-GT insertion, successfully generating the RosaBxb-GT/GA allele in B6 and NSG; see **Table 1**. After a minimum of one back-cross to the appropriate strain background these were bred to homozygosity; see **Table 2**.

### Use of dual site RosaBxb-GT/GA strains to generate RMCE Alleles

Typically, homozygous males carrying the dual landing pad are mated to super-ovulated wild-type females, generating heterozygous embryos for microinjection with Bxb1-Int mRNA and various DNA donor plasmids. *Highly* purified donor plasmid carrying dual cognate attB sites flanking the donor transgene were microinjected into zygotes as outlined in **Fig. 5**, and results summarized in **Table 4**. Twenty-five unique transgenic alleles were created at efficiencies ranging from 3 to 43%, with a combined average efficiency across all projects of 15% RMCE positive alleles, with constructs ranging in size from 1.5 kb to ∼43 kb. As highlighted in **Table 4**, a few projects initially performed poorly (P-7, P-12, P-15 and P21). However, after repeating the microinjection with a newly purified donor DNA, these generally succeeded, often with dramatically improved results.

**Fig. 5.**
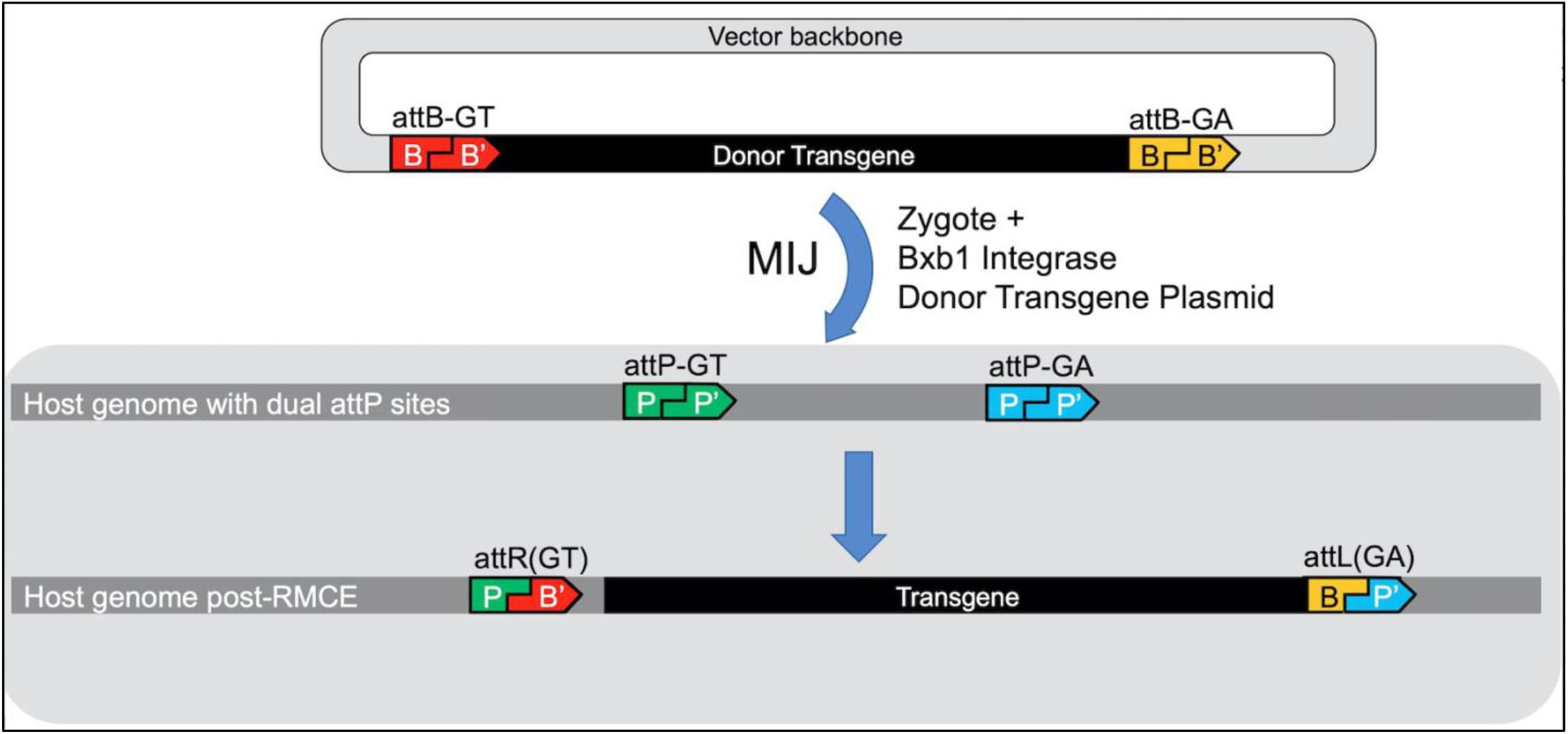
Schema for the use of dual attP-GT/attP-GA mouse strains targeted for transgenesis by Recombinase-Mediated Cassette Exchange (RMCE). Bxb1 integrase mRNA, the vector carrying the donor transgene (flanked by dual attB-GT, attB-GA sites) are microinjected into zygotes carrying the cognate attP-GT, attP-GA sites. The RMCE excludes the vector backbone and results in the precise integration of the desired donor transgene into the ROSA26 locus.

**TABLE 4:** Summary of RMCE attempts. **Successful Targeted Transgenesis in Dual Site RosaBxb-GT/GA Strains** Summary of 29 attempts to integrate 25 unique donor transgenes into the RosaBxb-GT/GA dual-site allele in B6 and/or NSG genetic backgrounds. Shown are DNA donor size, the number of microinjected embryos transferred, and the number of positive Founders expressed as a percentage of the number of animals screened. The Krt18-ACE2 construct (P-11) was delivered to both B6.RosaBxb-GT/GA and NSG.RosaBxb-GT/GA hosts. For a few constructs (P-7, P12, P-15, P-21), the microinjection was repeated following re-purification of the donor vector resulting in an improved RMCE rate.

| B6 Background (B6.RosaBxb-GT/GA) |  |  |  |  |
| --- | --- | --- | --- | --- |
| Donor ID | Donor Size (kb) | # embryos | RMCE POS | Note |
| P-1 | 1.5 | 78 | 9/23 (39%) |  |
| P-2 | 3.2 | 128 | 1/40 (3%) |  |
| P-3 | 3.3 | 130 | 2/36 (6%) |  |
| P-4 | 3.3 | 130 | 5/36 (14%) |  |
| P-5 | 3.7 | 129 | 16/37 (43%) |  |
| P-6 | 3.7 | 149 | 8/48 (17%) |  |
| P-7 | 4.5 | 63 | 0/11 (0%) | Rescued by repeat! |
| P-7 (RPT) | 4.5 | 106 | 3/22 (14%) |  |
| P-8 | 4.6 | 158 | 10/43 (23%) |  |
| P-9 | 5.7 | 165 | 6/50 (12%) |  |
| P-10 | 5.8 | 140 | 7/39 (18%) |  |
| P-11 | 6.8 | 182 | 10/54 (19%) | Krt18-ACE2 |
| P-12 | 7.1 | 150 | 1/31 (3%) | Improved by repeat! |
| P-12 (RPT) | 7.1 | 190 | 12/63 (19%) |  |
| P-13 | 8.5 | 105 | 1/33 (3%) |  |
| P-14 | 8.8 | 62 | 1/14 (7%) |  |
| P-15 | 9.4 | 73 | 0/22 (0%) | Rescued by repeat! |
| P-15 (RPT) | 9.4 | 143 | 8/44 (18%) |  |
| P-16 | 10.3 | 67 | 2/11 (18%) |  |
| P-17 | 11.5 | 102 | 1/37 (3%) |  |
| P-18 | 12.3 | 137 | 4/36 (11%) |  |
| P-19 | 13.0 | 115 | 2/21 (10%) |  |
| P-20 | 13.0 | 104 | 1/33 (3%) |  |
| P-21 | 23.6 | 88 | 0/28 (0%) | Rescued by repeat! |
| P-21 (RPT) | 23.6 | 107 | 2/26 (8%) |  |
| P-22 | 25.9 | 112 | 2/26 (8%) |  |
| P-23 | 30.6 | 139 | 4/35 (11%) |  |
| P-24 | 42.9 | 129 | 3/21 (14%) |  |
| **TOTAL** | **1.5-42.9kb** | **3381** | **121/890 (14%)** |  |

| NSG Background (NSG.RosaBxb-GT/GA) |  |  |  |  |
| --- | --- | --- | --- | --- |
| Donor ID | Donor Size (kb) | # embryos | RMCE POS | Note |
| P-11 | 6.8 | 260 | 18/63 (29%) | Krt18-ACE2 |
| P-25 | 9.1 | 162 | 8/20 (40%) |  |
| **TOTAL** | **6.8-9.1kb** | **422** | **26/83 (31%)** |  |

**Table 4: Successful Targeted Transgenesis in Dual Site RosaBxb-GT/GA Strains**
| All Backgrounds Combined (B6 & NSG) |  |  |  |  |
| --- | --- | --- | --- | --- |
|  | Donor Size (kb) | # embryos | RMCE POS | Note |
| **TOTAL** | **1.5-42.9kb** | **3803** | **147/973 (15%)** |  |

Founder offspring carrying the donor insertion were backcrossed to their respective genetic background generating N1 animals for germline transmission and complete validation of the transgenic allele. Screening was performed using a combination of PCR strategies and Sanger sequencing of product as described. A limited number of alleles were subject to nCATS to completely verify the precise nature of the transgene in its genomic context. On rare occasions (>1%), we did identify instances where either a mis-matched recombination event occurred (i.e., attB-GA + attP-GT) or only one of the two sites successfully recombined. It should be emphasized that these were rare events, see (55), and these alleles were easily identified by our screening strategy allowing for the aberrant lines to be detected and discarded. We did detect off-target, random integrations of the donor DNA at expected frequencies for random transgenesis (∼5% or less). Typically, Founders carrying an off-target allele were discarded. On the rare occasion where a positive founder also carried a random integration these, mice were backcrossed to segregate and select for the correct integration event.

### The rapid development of a Krt18 driven human ACE2 transgenic model in B6 and NSG backgrounds

Based on a construct used to make the random transgenic mouse line B6.Cg-Tg(K18-ACE2)2Prlmn/J JAX# 034860 (56), a near identical 6.84kb construct (“P-11”) was assembled to express human ACE2 under the direction of a human Krt18 promoter. Using Bxb1 Int-mediated transgenesis this construct was introduced into the B6.RosaBxb1-GT/GA and NSG.RosaBxb1-GT/GA strains. Initial characterization of offspring detected the correctly targeted transgene in Founder animals in the B6 background at 19% (10/54) and in the NSG background, 29% (18/63) – these data are summarized in **Table 4**. Carrier Founders were backcross to their respective background and confirmed faithful germline transmission. The resulting heterozygous targeted offspring were identified and the transgene confirmed by PCR and Sanger sequencing. These N1 animals were determined to be free of random transgenes and were also verified by nCATS. For both strains, heterozygous animals were available for experiments ∼4 months from the date of injection. Both lines were bred to homozygosity and deposited at the repository of the Jackson Laboratory (see **Table 2**).

### Comparative transgene sequence verification of the B6.RMCE[Krt18-ACE2] allele versus a Random ACE2 transgenic strain B6.Cg-Tg(K18-ACE2)2Prlmn/J

The development of a targeted transgenesis strategy was driven by the ever-increasing need for efficient and precise genetic engineering of animals. To fully verify insertions of increasingly larger DNA constructs we developed a DNA Nanopore sequencing workflow, based on nCATS, which provides enrichment of reads spanning the transgenic insertion (54). Currently, we have applied this targeted sequencing workflow to Bxb1-int mediated transgenic strains with insertions of 5 to 43 kb. In **Fig. 6a** we also present an example of the nCATS workflow and the resulting characterization of the B6.RMCE[K18-HuACE2] mouse. This allele was validated by gRNA targeting and subsequent enrichment of an 8.5kb region, generating an average per-base coverage of 195X over the transgene and surrounding chromosomal region (**Fig. 6b**). These nCATS-derived data, which include 800bp and 900bp flanking genomic (ROSA26) sequence, fully support that the transgenic sequence integrated precisely as expected.

**Fig. 6.**
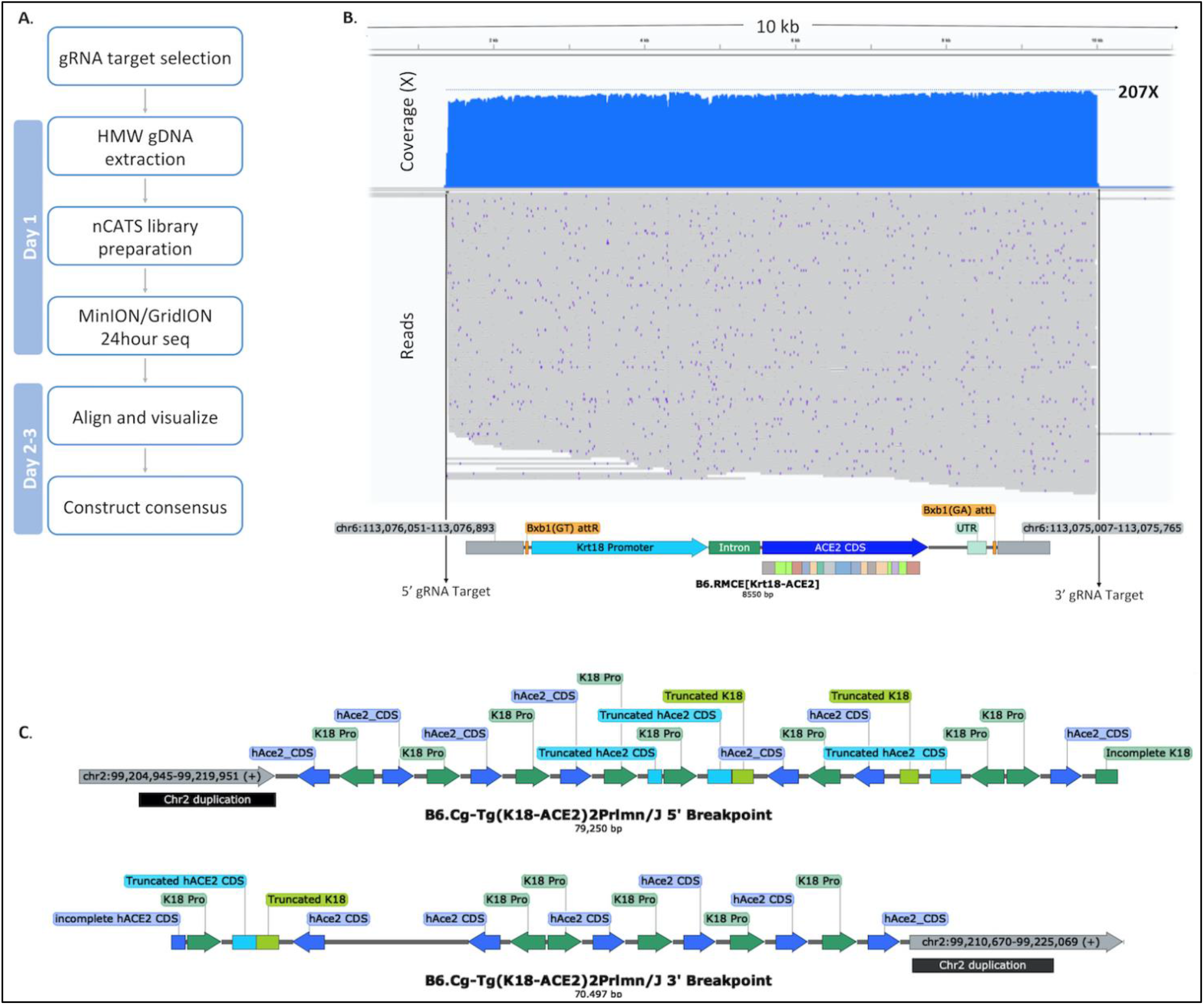
Characterization of transgenic mice using nCATS. A) Post sequencing workflow to construct insert regions and identify genomic locations. B) nCATS characterization of B6.RMCE(Krt18-ACE2) allele generated using the Bxb1 Int to achieve single copy integration, 215 reads on-target generating 195X coverage across the entire 8.5kb insertion site and 900/800bp genomic breakpoints. C) nCATS characterization B6.Cg-Tg(K18-ACE2)2Prlmn allele, 5’ and 3’ breakpoints identified at least 11 copies of K18-ACE2 in varying orientations.

To illustrate the precision of Bxb1 Int-mediated transgene integration compared to conventional random integration, we applied nCATS to the B6.Cg-Tg(K18-ACE2)2Prlmn/J mouse (JAX Stock #034860). Guides (**Supplemental Table 1**) targeting the suspected integration site on Chromosome 2 (chr2:99204917-99225067) as well as against the CDS of ACE2 revealed the complex architectural nature of this random transgenic insertion. These data lead to a reconstruction of a 79.3 kb 5’ and 70.5 kb 3’ breakpoint sequences (**Fig. 6c**) and identified K18-ACE2 concatemers consisting of at least 11 copies in both orientations. In addition, we also detected a 10.5kb duplication of Chromosome 2 at the periphery of each breakpoint along with K18-ACE2 cassette truncations at sites of inversion throughout the insertion. Due to the repeat-rich complexity of the insertion we were unable to generate a full consensus for the entire integration site.

## Discussion

There is an essential need to create genetically modified animals where large transgenes are integrated precisely, in a defined genomic context to further advance biological research and transgenic development (3, 6, 57). Although targeted nucleases can accomplish this to a degree, their efficiency in the placement by HDR of donor constructs beyond a few kb is unpredictable and in general too limited. To address aspects of this we developed a strategy enabling efficient precision integration of large DNA constructs into the ROSA6 safe harbor locus directly via the zygote across multiple mouse genetic backgrounds. To execute our strategy we required high efficiency, site-specific recombination which is not significantly impacted by donor constructs of up to at least 10 kb. For this we selected from the limited number of lysogenic bacteriophages tested for targeted integration into the mammalian genome, Bxb1 integrase. This integrase has proven specificity, lacks pseudosites in the mouse genome and is reported to have the highest activity in cell lines when compared with other serine integrases, including φC31 (46, 55, 58). Further, as the bacteriophage Bxb1 has a genome size of 50.5kb we hypothesized that this site specific recombinase could recombine transgenes to at least this size.

### CRISPR/Cas9 introduction and use of a single Bxb1 attP-GT site

To test the Bxb1 Int approach, we used CRISPR/Cas9-mediated insertion of the Bxb1 attP site with the wild type “GT” central dinucleotide, i.e., attP-GT, into the ROSA26 locus. This was done in three commonly used inbred mouse backgrounds, B6, FVB and the immunocompromised line NSG to help assess its capabilities. Initially, for C57BL/6J we used a CRISPR/Cas9 NHEJ mediated dsDNA oligo strategy based on GUIDE-seq (53). Although this approach worked, the efficiency was low (7% carriers) and also led to damage at the integration site. For subsequent work we used CRISPR/Cas9 ssDNA oligo mediated HDR into the same site in ROSA26 with mouse strains NSG (73-100% carriers) and FVB (19% carriers) (see **Fig. 3, Table 1**).

Once established, we used these modified strains (B6.RosaBxb1-GT, FVB.RosaBxb1-GT and NSG.RosaBxb1-GT) to test Bxb1 Int-mediated integration focusing on establishing MIJ conditions, including Bxb1-integrase mRNA concentration and minicircle DNA isolation (see **Fig. 4**). To aid in candidate identification, we developed a series of rapid PCR screening and sequencing strategies which included the flanking regions of the ROSA26 integration attP-GT site. This allowed complete sequence verification of Bxb1 Int-mediated insertions in the context of the ROSA26 locus. Our tests, summarized in **Table 3**, show clearly that targeted integration into ROSA26 attP-GT occurs at levels at or more often, considerably higher than random transgenesis, ranging from 1 to 33% with constructs of 3.2 to 15.3kb.

### Development of a genetically diverse panel of RosaBxb1-GT strains

A key driver of this work was to develop improved access to efficient transgenic targeting using multiple genetic backgrounds, facilitating comparative genetic background studies. Our success in construction of and using the ROSA26 Bxb1-GT allele in three backgrounds inspired us to add the same attP site/location to a host of novel genetic backgrounds. The choice of these focused on the Collaborative Cross founders and also included DBA/2J due to its importance in developing the BXD RI strains (59, 60).

The genetic modification of many genetic backgrounds has been hindered as many of these “non-conventional” mouse strains exhibit poor zygote yield and/or survivability post modification attempts. Although mouse embryonic stem cell (mESC) lines exist for many backgrounds and could be used to circumvent this, the introduction of constructs into these is often encumbered by drug selection markers, requires additional time to produce the chimeras and (possibly) obtain germline transmission, as compared to direct modification of zygotes from the required background (61-63). Also, by using extant mouse strains the actual genetic background of the resulting models will represent the present genome of the background and not that of a long-established mESC line (64).

Although CRISPR/Cas9 mediated HDR with ssDNA oligonucleotides is relatively efficient, embryo yield and survivability from some genetic background are problematic. Our initial attempts to introduce the attP-GT site used microinjetion of zygotes. However as shown in **Table 1**, the results were mixed on both success to live born and introduction of the attP-GT site into the ROSA26 locus. This difficulty was especially evident in the wild inbred strains CAST, WSB and PWK, which failed completely due to a low embryo yield/female, compounded by subsequent poor survivability to term following MIJ. The use of electroporation as an approach to introduce CRISPR/Cas9 and *small* ssDNA donors has been shown by others to be efficient in B6 mice (51, 52). Using conditions previously established for B6 zygote electroporation we introduced the attP-GT site into a series of different genetic backgrounds. Although multiple direct comparative experiments of the overall efficiency of MIJ vs electroporation were not part of this study, we did succeed in obtaining attP-GT site integration in multiple strains, including CAST and PWK (**Table 1**), using electroporation. A review of these data suggests that the success is linked to the higher survivability of zygotes to term post electroporation as compared to microinjection, although further experiments would be needed to fully confirm this hypothesis. Combined, these efforts generated a diverse genetic panel of ten strains (listed in **Tables 1 and 2**) carrying the attP-GT site in the ROSA26 locus. However, even with our best efforts we failed at this time to generate the desired allele in the wild inbred mouse strain WSB, due to poor availability of zygotes combined with low survivability post electroporation.

Out of concern for potential off-target integration of the donor attP-GT sequence, we modulated the concentration of the oligonucleotide delivered by EP in some of our tests. Although we did not screen for off-target integration, we did find that greatly reducing the level of donor oligo in the electroporation does not necessarily result in reduced knock-in efficiency. For example, despite reducing the concentration of the donor by more than 90% (to 70 ng/µl) when targeted NOD/ShiLtJ embryos, we still successfully generated the intended allele with 57% efficiency (see **Table 1**). We believe that this parameter should be further investigated.

### Development and use of dual Bxb1-att sites

Precise Bxb1 Int-mediated integration of donor constructs into the ROSA attP-GT site was demonstrated in three genetic backgrounds. However, we sought to further improve this by exploring the use of a second Bxb1 attachment site (**Fig. 3**). We reasoned that the second site in close proximity to the initial attP-GT site would increase stochastic interaction and hence integration. A key attribute of using a dual heterologous attachment site strategy is that the donor vector can now be designed to integrate by RMCE, leading to excision and loss of the plasmid backbone and directional integration of the construct (see **Fig. 5**) (65). We selected the GA dinucleotide for our second site based on Jusiak et al’s work (55) which compared wild type Bxb1-GT to the Bxb1-GA variant in cell lines and noted that the GA variant resulted in more than double the integration efficiency. They also reported minimal crosstalk between Bxb1 GA and GT att sites, an essential requirement when using dual heterologous sites (55). We added the Bxb1 attP-GA site *in cis* to the attP-GT site in B6.RosaBxb-GT and NSG.RosaBxb-GT mouse strains by CRISPR/Cas9 mediated ssDNA oligo HDR, creating the strains B6.RosaBxb-GT/GA and NSG.RosaBxb-GT/GA respectively. Using these lines tests were conducted with a range of constructs (See **Fig. 5**), and data summarized in **Table 4**. This process improvement also lead to operational advantages, including avoiding troublesome minicircle preparation, use and artifacts. Surprisingly, when using these dual attP site mice with cognate constructs we saw a more than three-fold increase in overall efficiency over similar tests with animals carrying a single attP-GT site. This increased efficiency is intriguing, perhaps reflecting synergy or simply an additive effect of the combined Bxb1-GT and the more efficient Bxb1-GA sites. RMCE may also improve operational efficiency, especially as transgene plasmid preparation is considerably simpler and potentially of higher quality (65-69). The use of RMCE also opens the possibility of constructs of tens, if not hundreds of kilobases, which would be difficult to prepare as minicircles at the concentration and purity required.

Our data is insufficient to ascertain if there is a strong correlation between construct size and efficiency of integration. However, it would be logical to assume that increasing the size of the insert is likely to decrease the integration efficiency. Using identical DNA concentrations for microinjection (e.g., 30 ng/µl), larger constructs are actually delivered at a lower *copy* number per zygote, and the reduced molar concentration of donor attB sites would be expected to impact integration efficiency due to a simple reduction in stochastic interactions (70). It also should be considered that larger constructs are more susceptible to DNA breakage during handling, including during MIJ, which would be expected to reduce their integration efficiency. To counter this, it may be worthwhile to incorporate some of the strategies previously described for BAC transgenics, notably modulating the MIJ buffer (71, 72). Our preliminary work on this front shows promise (data not shown), however an increase in aberrant insertion events suggests the need for greater optimization.

### Speed of development example - Krt18-ACE2 transgenic in two genetic backgrounds

To demonstrate the Bxb1-int approach’s speed and to aid SARS-CoV research, two novel mouse models were created based on a previous random Krt18 driven human ACE2 transgenic mouse line (56). Using dual aatP sites we produced a large number of ROSA26 targeted transgenics enabling the development of sequence-verified, single copy, N1 heterozygous transgenic alleles in the B6 and NSG backgrounds in under five months. Bxb1-int mediated transgenesis is directional, for the Krt18-ACE2 lines developed here the transgene was inserted in the same orientation as Gt(Rosa26)Sor lncRNA. Ideas concerning which direction is optimal for transcription of inserted transgenes into the ROSA26 remain unresolved, and are undoubtedly complex involving the particular integration site and the construct’s promoter (32, 34). Preliminary work on these two strains indicated that the ROSA26 transgene human ACE2 under the Krt18 promoter is transcribed at high levels in both B6 and NSG backgrounds (data not shown).

### Complete transgenic sequence verification and comparison to random transgenesis

A central tenet of the approach outlined here is to provide precision and definition of DNA integration. In pursuit we also developed methods to deliver complete sequencing of integrated DNAs. An approach coupled with recent developments in amplification-free targeted long read sequencing by nCATS used the Oxford Nanopore minION/GirdION platform. This allowed for example here, validation of >43kb insertions including ∼1kb of the flanking regions. As expected, nCATS verification of the Bxb1 Int-generated Krt18-ACE2 strains showed a single on-target integration with no disruption to the surrounding genomic sequence. These data demonstrate that the Bxb1 Int system can precisely integrate transgenes intact into a known genomic location and this can be rapidly validated including the surrounding genomic region. Combining the Bxb1 system with nCATS as demonstrated here, allows us to construct models with ever increasing confidence with reduced/no host chromosomal deformation.

In contrast, when nCATS was used to characterize the random transgenic B6.Cg-Tg(K18-ACE2)2Prlmn/J mouse we clearly saw the complexity that traditional random integration transgenesis can lead to. Fortunately in this case the random Krt18-ACE2 transgene integration did not lead to disruption of any annotated genes. However, it illustrates how random integration can adversely impact the host genomic environment, as shown here we identified a partial duplication of Chromosome 2 (**Fig 6c**). Additional models generated via random integration transgenesis have been assayed with nCATS, and every model assayed showed concatemerization and often deletion of large sections of the host chromosome. We have also observed the co-integration of the plasmid backbone and large 80 kb+ BAC fragments of donor DNAs (data not shown).

Lastly, often upon comparison with reference sequences, we notice that these and other unpublished data can show that integrated construct sequences, especially those derived from genomic libraries, differ at occasional bases. These differences were also confirmed when sequencing the originating microinjected donor DNA (data not shown). In general, these polymorphisms appear to be silent or (appeared) irrelevant to the overall construct’s function. However, this may not always be the case. As such, complete sequence characterization of the integrated constructs provides an elevated level of confidence in the genetic modification achieved and the resulting phenotype.

### Conclusion and future directions

The strategy devised and presented here was envisioned to be used in any laboratory with access to zygote microinjection and general assisted reproductive technologies. The overall operational advantages include: scalability, high efficiency, rapid screening and defined precision integration of constructs to at least 43kb. The approach can be used on multiple genetic backgrounds extending ideas of genetic diversity and crucially providing a uniform integration site for expression of transgenes. The attP/B approach also makes feasible ideas of subsequent sequential gene addition or “gene-stacking” by embedding other heterologous att sites in introduced constructs or by multipart assembly in vivo (44, 73, 74). The advent of advanced sequencing protocol/methods exemplified here in the single molecule reading of ∼80kb further advances the reality of precision in the genetic modification of animals. Future directions with site specific recombinases include the potential to isolate novel integrases and/or manipulate targeting via directed evolution, enhancing efficiency through integrase site engineering, reprogramming specificity, or fusion to, e.g., Cas9+gRNA, enabling targeting insertions to specific naturally occurring sequences within the genome (39, 41, 48, 75, 76). Such designer site specific recombinases would have use in synthetic biology, biotechnology and genetic engineering, including human gene therapy.

## Acknowledgments

We thank Cindy Avery and Todd Nason for excellent maintenance of mouse colonies, Richard S. Maser for building the Krt18-ACE2 construct, and Peter Kutny and group at Genetic Engineering Technologies for microinjections. We also thank various groups, including the Chakravarti laboratory (Dr Ryan Fine at NYU School of Medicine), the Reinholdt laboratory (Dr David Bergstrom and Tiffany Leidy-Davis at JAX), and the Steve Murray laboratory (Dr. Kathy Snow at JAX) for providing the constructs. We are grateful to the Genome Technologies (GT) group at JAX for their Sanger and Nanopore sequencing services. We thank Melissa Berry for her careful reading of the manuscript and assistance in generating the mouse strain nomenclature. This work was funded by National Institutes of Health grants P30CA034196, R21 OD027052, and R24 OD 021325 (Drs. Reinholdt and Bergstrom). The Krt18-ACE2 related work was partially funded by the Rosenthal laboratory at JAX. We also gratefully acknowledge funding from JAX.

## Ethics Statement

The Institutional Animal Care and Use Committee of The Jackson Laboratory approved all procedures used in this study and all mice were maintained at The Jackson Laboratory (Bar Harbor, ME, USA) in strict accordance with all institutional protocols and the Guide for the Care and Use of Laboratory Animals.

## Author contributions

B.E.L., V.H., and M.V.W. conceptualized the project. B.E.L., V.H., and M.V.W. designed the experiments. B.E.L prepared the reagents for microinjections and performed the DNA gel electrophoresis experiments. S.L. developed and prepared the reagents for nCATS and performed the experiments. B.E.L., S.L., V.H., and M.V.W. analyzed the data and wrote the manuscript. All the authors contributed to data interpretation, critical review, and approval of the manuscript.

**Supplemental Table 1.**
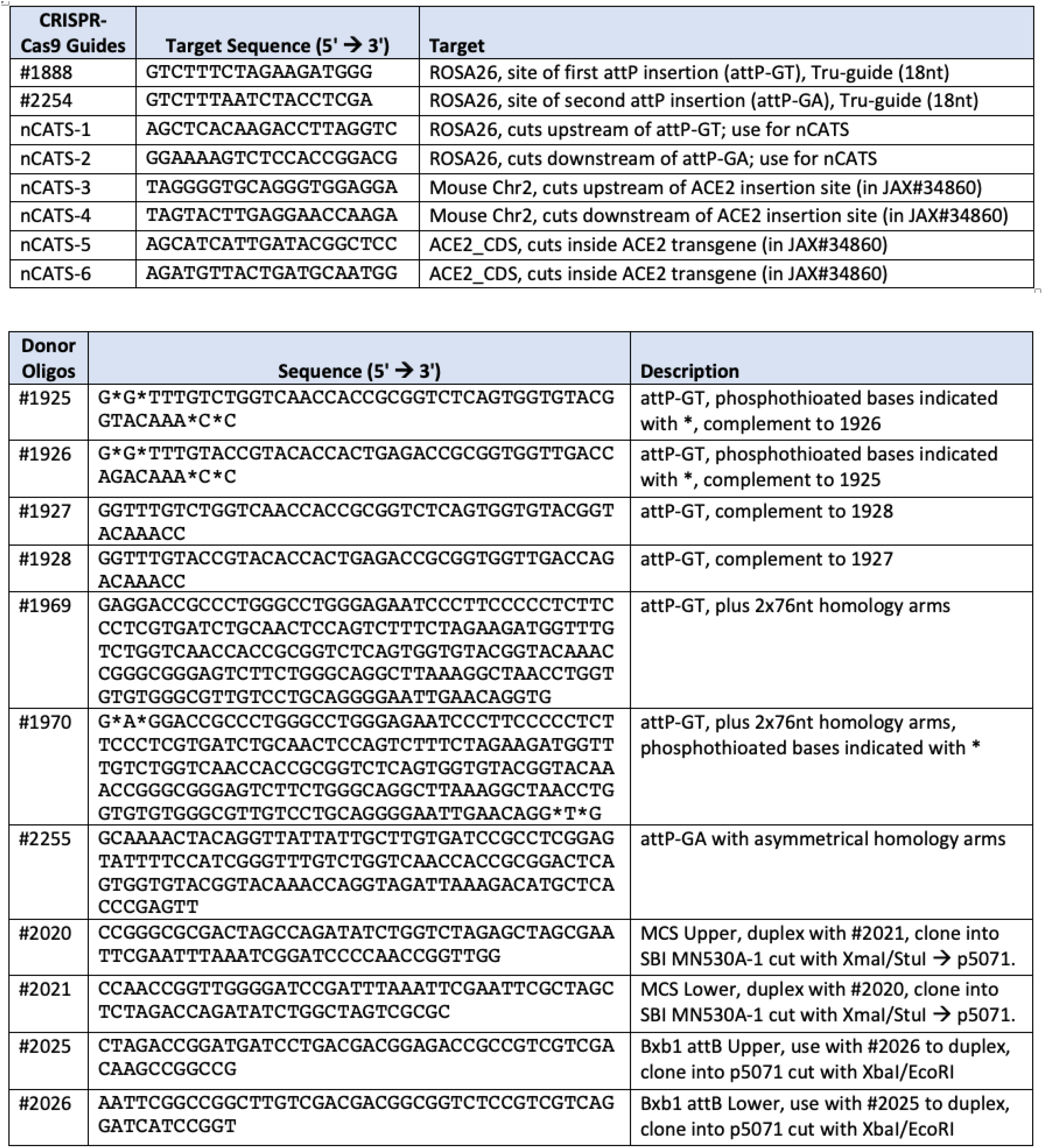
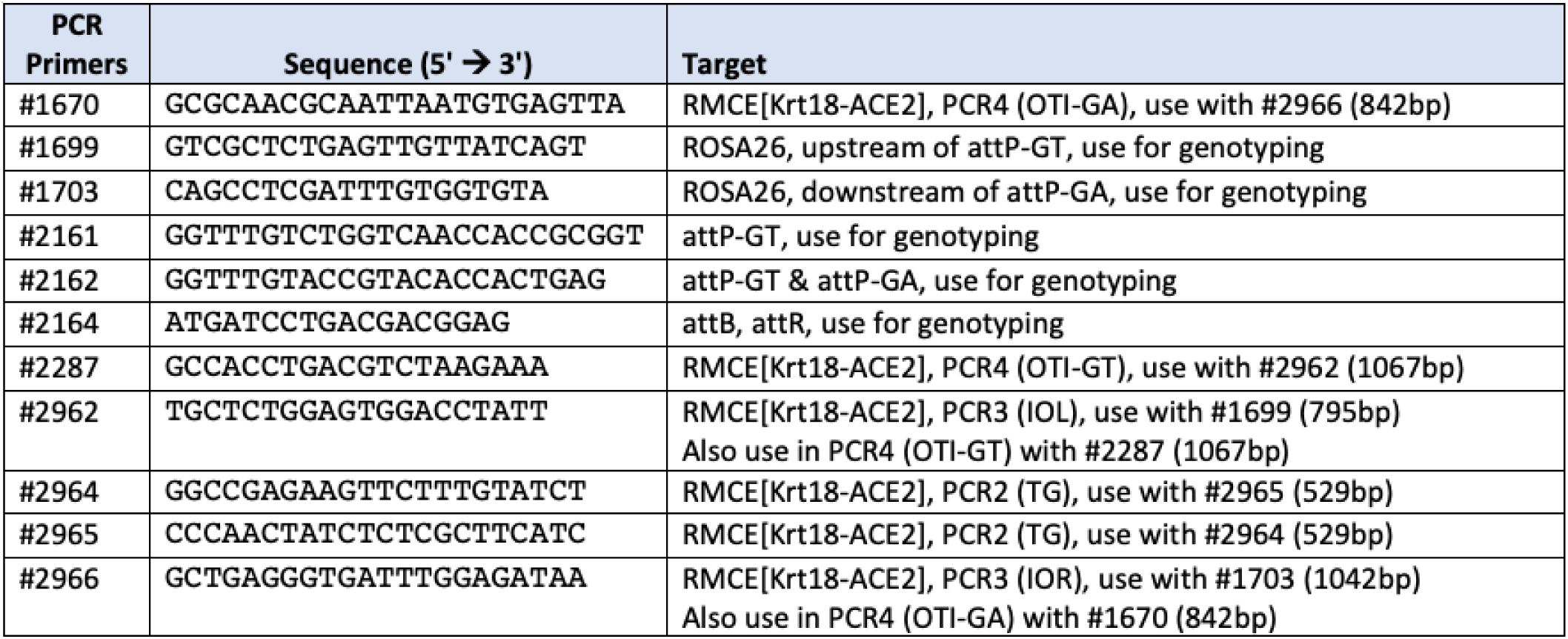

